# From Peaks to Patterns

**DOI:** 10.1101/2025.11.09.687456

**Authors:** Hossein Fathollahian, Marziye Salahshour

## Abstract

MRS is a non-invasive method for gathering information about cellular metabolism and can be used to study neurological and metabolic disorders. However, the clinical utility is limited by low signal quality, overlapping spectral peaks, and the inability to interpret and compare metabolite data. We developed an interactive visual analysis system that integrates methods for filtering, quality measurement, and comparative summaries of MRS signals, enabling researchers to more effectively evaluate and interpret spectral data. This will enable researchers to conduct a more accurate evaluation of spectral data, thereby facilitating a deeper understanding of the molecular processes underlying complex neurological and metabolic processes.

## 1 Introduction

The non-invasive nature of ^31^P-MRS lends itself to examining the metabolic activity of the brain, which reflects the underlying biochemical changes that may not be observable until the disease is structurally or histopathologically affected. This is uniquely valuable for studying neurodegenerative diseases. Nonetheless, the problems of intrinsic noise, frequency misalignment, and spectral complexity make it challenging to identify the metabolites correctly, estimate quality, and compare them. We address the problems of noise, misalignment, and spectral complexity by exploiting sophisticated wavelet-based denoising and accurate frequency-shift correction, thereby improving spectral fidelity and alignment. Following these advances, we present an interactive visualization system that utilizes newly developed circular and rectangular encoding, allowing the ratios of metabolites and the quality of spectra to be browsed in flexible views that can be more reliably explored to create more holistic characterizations of metabolic patterns.

**Figure 1.**
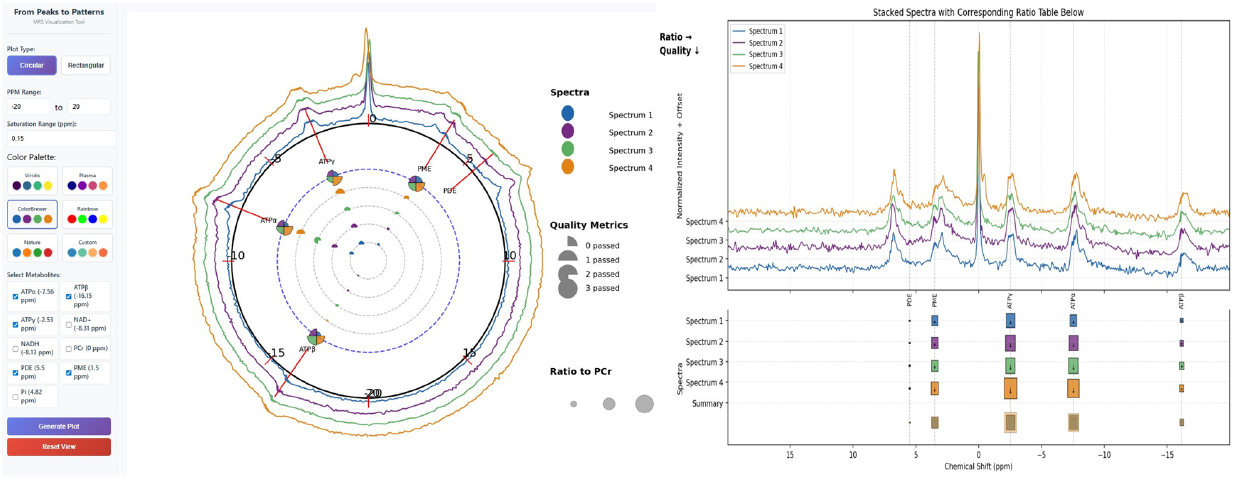
Dual MRS Visualization Interface. (A) Metabolite selection panel; (B) Circular plot showing ratio (radius) and quality (saturation) with summary slices; (C) Rectangular plot displaying ratio (width), quality (height), and summary heatmap.

## 2 Related Work

MRS data are pre-processed to eliminate noise and artifacts, then this processed data is quantified to eliminate the effects of low SNR and peak overlaps with spectral integration or model fitting [3, 2, 6]. Interactive functionality, including ComVis+MITK [4], Polaris [5], and SpectraMosaic [1], is also useful, especially the ratio analysis with user-defined glyphs and with a heatmap in cohort-level analysis with SpectraMosaic. Nonetheless, the systems typically lack a unified visualization of metabolite ratios and the quality of the MRS signal. Instead, we leverage these strengths to create a new glyph-based visualization that adds more informative new features, such as ratio summarization, quality measurement, and integrated views that improve the perception of spectral quality and peak separation in combination with comparative analysis.

## 3 Redesign

Within the redesigned pipeline (Fig.2) every spectrum *S* ∈ ℝ ^2048^ is denoised using a soft-thresholded version of the discrete wavelet transform (Daubechies-4) to eliminate high frequency noise but manage to retain peaks. The peaks are then identified through the local maxima algorithm with amplitude and prominence constraints, and spectra are aligned by frequency using the phosphocreatine (PCr) reference peak. The metabolite trends will be quantified with the integrals of preselected spectral windows and normalized to associated PCr, and will be evaluated by the Full Width at Half Maximum (FWHM), Cramér–Rao Lower Bound (CRLB), and SNR [2]. The aligned peak-detected and denoised spectra are demonstrated in Fig.3, and the end result is presented in circular and rectangular form designs. In the circular glyph-based representation (Fig. 4), each metabolite is positioned along an orbit, with its distance from the center (radius) encoding the metabolite-to-PCr ratio. Each circle is divided into four wedge-shaped quadrants to represent peak quality: one filled quadrant indicates failure of all quality measures, two quadrants indicate passing only one, three indicate passing two, and all four quadrants indicate passing all measures. The saturation of the circle encodes peak area. Finally, the outermost orbit summarizes the average metabolite ratios across all spectra, using the same radius encoding for consistency.

**Figure 2.**
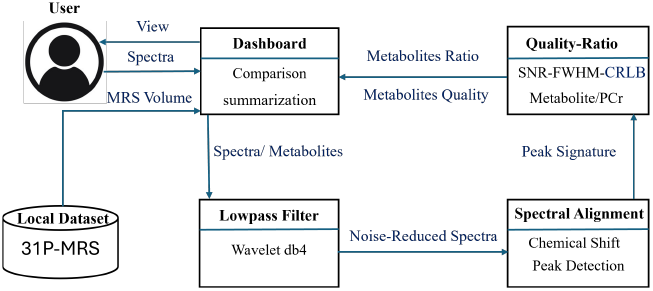
Workflow illustrating metabolite peak detection, denoising, ratio/quality extraction, and visual comparison.

**Figure 3.**
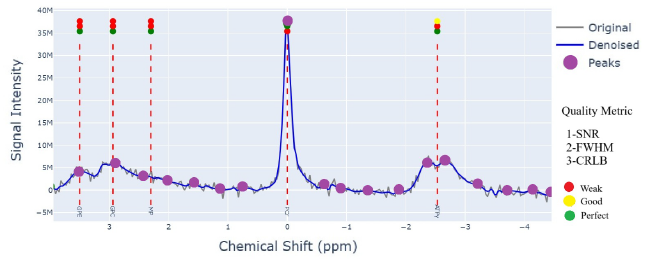
Original (black) and denoised (blue) spectra with metabolite lines (red), Local peak (green), and quality metrics(traffic light).

**Figure 4.**
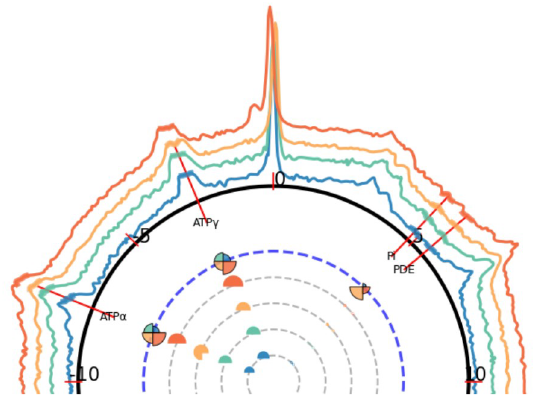
Circular MRS plot with circles showing ratio to PCr (radius), quality (fill), and peak area (saturation), with outer orbit summarizing all spectra.

**Figure 5.**
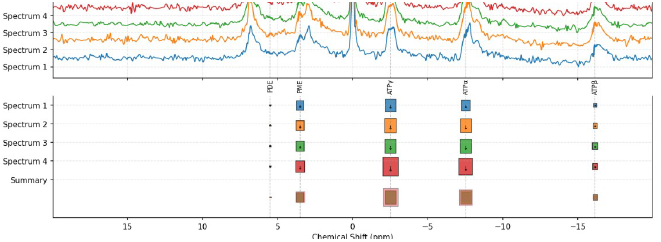
Rectangular MRS plot with bars showing ratio to PCr (length), quality (width), and peak area (saturation), The lowest row provides a combined summary of all spectra for comparison.

On the rectangular view, spectra are organized into rows; the rectangle’s length represents the metabolite-to-PCr ratio, and the width reflects quality. The component of the total spectra has a summary overlay, which gives a ratio distribution along the length and the quality distribution along the width, which offers a parallel perspective of the metabolite ratios to PCr and peak quality. Reliable metabolite-to-PCr ratios from peak areas are visualized in circular and rectangular layouts for comparison across ppm Figure 4: Circular MRS plot with circles showing ratio to PCr (radius), quality (fill), and peak area (saturation), with outer orbit summarizing all spectra.

## 4 Result

The results are illustrated in both circular (Fig.4) and rectangular (Fig.5) displays, in which radius or length represents the metabolite-to-PCr ratio and fill saturation proportions to peak area to find out stronger signals. A standard color map is used, but the user may specify a custom-defined color map. In circular mode, individual spectra are emphasized by a summary orbit. In contrast, in rectangular mode, spectra are stacked and overall trends are indicated by an overlay, at which ratio and quality patterns are readily discerned.

## Supporting information

Supplemental output

## 5 Discussion and Conclusion

The reliability and interpretability of 31P-MRS are enhanced when PCr-based alignment combines with wavelet denoising. The proposed visualization can effectively depict metabolite concentration and quality, but the visual task (due to color limitations and signal overlap) may be complicated for human perception. PCr signals can interfere with alignment. Still, the Application also facilitates robust, comparative analysis and increases the availability of 31P- MRS in research and clinical fields. The development of visualization methods will be furthered, and biochemical

## References

[1] L. A. Garrison, J. Vasícek, R. Grüner, N. N. Smit, and S. Bruckner. Spectramosaic: An exploratory tool for the interactive visual analysis of magnetic resonance spectroscopy data. In VCBM, pp. 1–10, 2019. 1

[2] J. Near, R. Edden, C. J. Evans, R. Paquin, A. Harris, and P. Jezzard. Frequency and phase drift correction of magnetic resonance spectroscopy data by spectral registration in the time domain. Magnetic resonance in medicine, 73(1):44–50, 2015. 1

[3] J. Near, A. D. Harris, C. Juchem, R. Kreis, M. Marjańska, G. Öz, J. Slotboom, M. Wilson, and C. Gasparovic. Preprocessing, analysis and quantification in single-voxel magnetic resonance spectroscopy: experts’ consensus recommendations. NMR in Biomedicine, 34(5):e4257, 2021. 1

[4] M. Nunes, B. Rowland, M. Schlachter, S. Ken, K. Matkovic, A. Laprie, and K. Bühler. An integrated visual analysis system for fusing mr spectroscopy and multi-modal radiology imaging. In 2014 IEEE conference on visual analytics science and technology (VAST), pp. 53–62. IEEE, 2014. 1

[5] C. Stolte, D. Tang, and P. Hanrahan. Polaris: A system for query, analysis, and visualization of multidimensional relational databases. IEEE Transactions on visualization and computer graphics, 8(1):52–65, 2002. 1

[6] F. Torkamani-Azar et al. Video quality measurement based on 3-d. singular value decomposition. Journal of Visual Communication and Image Representation, 27:1–6, 2015. 1

