## Supplemental output for "From Peaks to Patterns"

Supplementary Material

### 1 Interactive MRS Discovery

We introduce an interactive framework for analyzing $^{31}$P-MRS spectra, integrating wavelet-based denoising and frequency-shift correction to address intrinsic noise, misalignment, and spectral complexity. Our system enables accurate metabolite identification, quality assessment, and comparative analysis through novel circular and rectangular visual encoding of metabolite ratios and spectral quality. Outputs are displayed online for scalable, expert-guided exploration of metabolic patterns in neurological and metabolic disorders. A live demonstration is available (`https://mrs-8w1w.onrender.com/`).

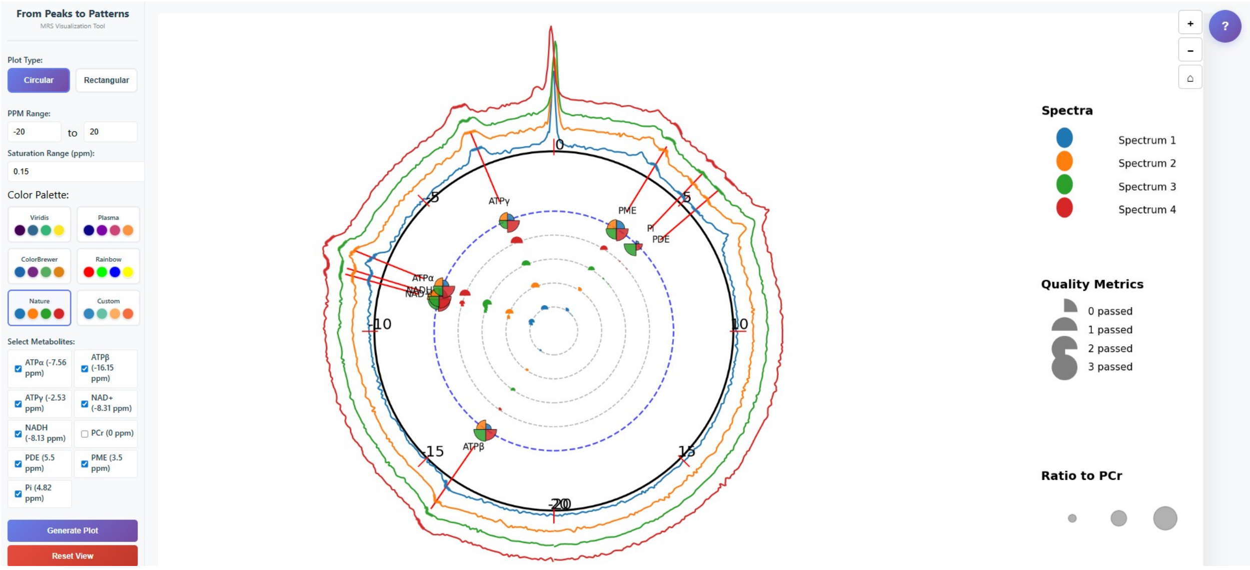


**Figure 1:** Circular visualization of $^{31}$P-MRS spectra using the "Nature" color palette. Colored traces represent four spectra, with metabolite markers where radius encodes ratio to PCr and filled segment proportion indicates quality.

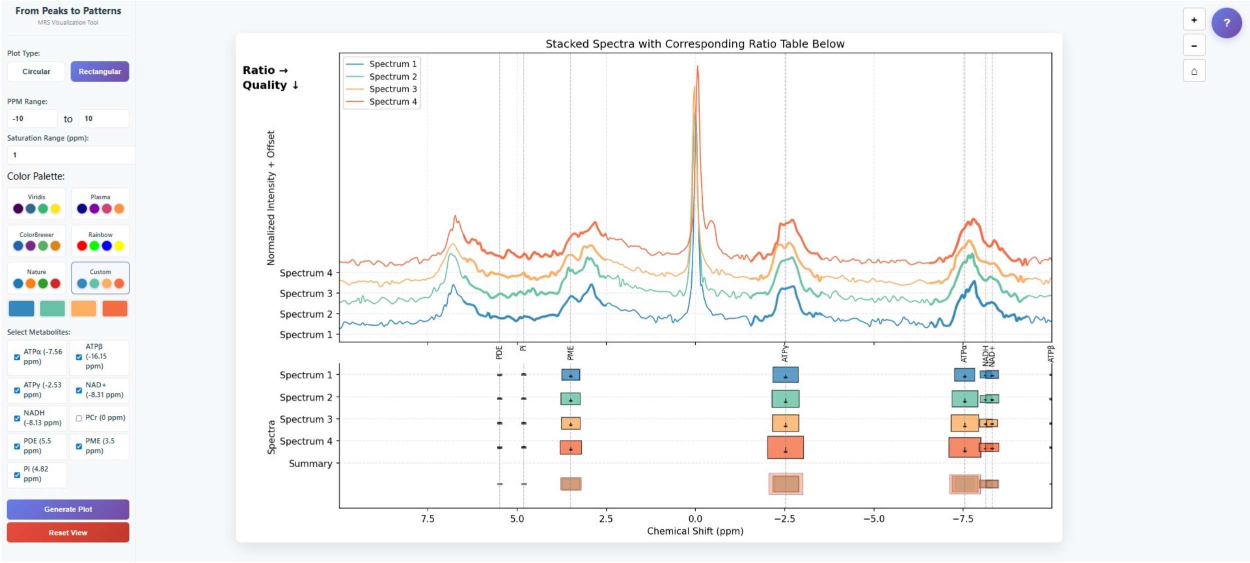


**Figure 2:** Rectangular visualization of $^{31}$P-MRS spectra using a custom color palette. Colored traces represent four spectra, with metabolite markers where bar length encodes the ratio to PCr and bar width indicates quality.

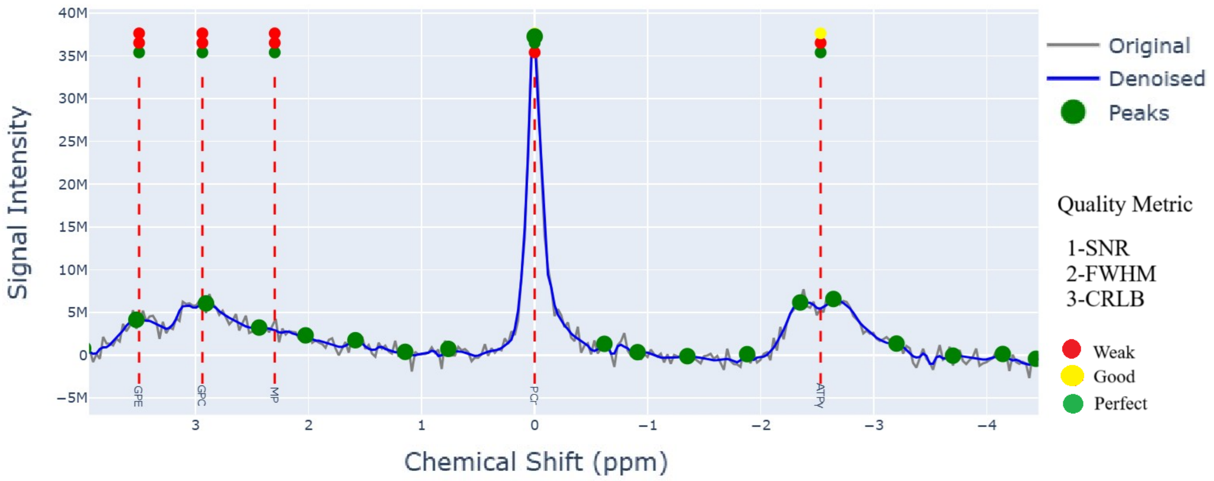


**Figure 3:** Denoised $^{31}$P-MRS spectrum with peak detection. Original (gray) and denoised (blue) spectra with detected peaks (green) and annotated metabolite positions (red dashed lines).
